# MHC Genetic Variation Influences both Olfactory Signals and Scent Discrimination in Ring-Tailed Lemurs

**DOI:** 10.1101/337105

**Authors:** Kathleen E. Grogan, Rachel L. Harris, Marylène Boulet, Christine M. Drea

## Abstract

Diversity at the Major Histocompatibility Complex (MHC) is critical to health and fitness, such that MHC genotype may predict an individual’s quality or compatibility as a competitor, ally, or mate. Moreover, because MHC products can influence the components of bodily secretions, an individual’s body odor may signal its MHC and influence partner identification or mate choice. To investigate MHC-based signaling and recipient sensitivity, we test for odor-gene covariance and behavioral discrimination of MHC diversity and pairwise dissimilarity, under the good genes and good fit paradigms, in a strepsirrhine primate, the ring-tailed lemur (*Lemur catta*). First, we coupled genotyping with gas chromatography-mass spectrometry to investigate if diversity of the MHC-DRB gene is signaled by the chemical diversity of lemur genital scent gland secretions. We also assessed if the chemical similarity between individuals correlated with their MHC similarity. Next, we assessed if lemurs discriminated this chemically encoded, genetic information in opposite-sex conspecifics. We found that both sexes signaled overall MHC diversity and pairwise MHC similarity via genital secretions, but in a sex- and season-dependent manner. Additionally, both sexes discriminated absolute and relative MHC-DRB diversity in the genital odors of opposite-sex conspecifics, supporting previous findings that lemur genital odors function as advertisement of genetic quality. In this species, genital odors provide honest information about an individual’s absolute and relative MHC quality. Complementing evidence in humans and Old World monkeys, our results suggest that reliance on scent signals to communicate MHC quality may be important across the primate lineage.

## INTRODUCTION

The Major Histocompatibility Complex (MHC) is an extremely polymorphic group of genes within the adaptive immune system of vertebrates that plays a critical role in disease resistance (Piertney & Oliver 2006). Because genetic diversity at the MHC is fundamentally linked to parasite resistance, survivorship, and reproductive success (Sommer 2005a, Piertney & Oliver 2006), an individual’s MHC genotype is hypothesized to be an important predictor of its quality as a mate. If recognizable to others, animals could increase their reproductive success by selecting mates that possess particular MHC genotypes, such as diverse alleles or specific disease-resistant alleles (Penn & Potts 1999, Mays & Hill 2004). Although researchers have found evidence that MHC genotype influences mate choice or its proxies in many species (reviewed in Kamiya et al. 2014), the mechanism by which animals assess the MHC of conspecifics is still under investigation (reviewed in Ruff et al. 2012). Given that the protein products of the MHC can influence body odor, scientists have implicated an olfactory-based mechanism (reviewed in Ziegler et al. 2005, Boehm & Zufall 2006); however, researchers rarely combine chemical and behavioral approaches within the same study to test the purported mechanism of information transfer (Leinders-Zufall et al. 2004, Milinski et al. 2005). Here, using the ring-tailed lemur (*Lemur catta*) – a strepsirrhine primate for which there is strong evidence of condition-dependent olfactory signaling (Charpentier et al. 2008, Boulet et al. 2010, Crawford et al. 2011) – we test for olfactory-based MHC advertisement and recognition. Specifically, we combine MHC genotyping with chemical analyses of genital secretions and with behavioral tests of scent discrimination between opposite-sex conspecifics to ask 1) if lemurs advertise their genetic quality and similarity at the MHC via chemical cues and 2) if conspecifics can detect this olfactory information.

Because an offspring’s health is influenced by the genotypes inherited from its parents, extreme polymorphism of the MHC is likely to be maintained both by health-mediated natural selection and by MHC-based sexual selection (Winternitz et al. 2013). Potential mates might be chosen for their MHC diversity, for their possession of a particular disease-resistant allele or for their MHC dissimilarity relative to the chooser (Sommer 2005a, Milinski 2014). Two models have been developed to explain the types of genetic benefits accrued owing to the potential partner’s genotype: In the ‘good genes’ model, individuals, regardless of their own genotype, choose ‘high-quality’ mates, with quality being defined by absolute or optimal genetic diversity or by possession of a particularly beneficial allele (Andersson 1994, Penn 2002, Neff & Pitcher 2005, Kempenaers 2007). This model is supported by research in several species of fish (Forsberg et al. 2007, Eizaguirre et al. 2009, Evans et al. 2012), and birds (Ekblom et al. 2004, Richardson et al. 2005).

In the ‘good fit’ model, individuals choose mates based on their relative genetic compatibility (Trivers 1972, Brown 1999, Neff & Pitcher 2005), so as to optimize parental genetic dissimilarity, minimize the risk of inbreeding or increase the genetic diversity of their offspring (Tregenza & Wedell 2000, Penn 2002). Support for this model (reviewed in Kamiya et al. 2014) derives from species of fish (Landry et al. 2001, Aeschlimann et al. 2003, Neff et al. 2008), reptiles (Olsson et al. 2003), birds (Bonneaud et al. 2006, Freeman-Gallant et al. 2003), and mammals (Radwan et al. 2008, Burger et al. 2015), including primates (Ober 1999, Schwensow et al. 2007 & 2008, Setchell et al. 2010, Kromer et al. 2016). The good genes and good fit models are not mutually exclusive, however, because a potential mate could be both maximally diverse and optimally dissimilar (Mays & Hill 2004). Individuals may also alter their preferences based on environmental conditions, current pathogen pressures or potential risk of inbreeding (Qvarnstrom 2001, Mays & Hill 2004).

Regardless of the model, for mate choice to be influenced by the MHC, individuals must both signal their respective MHC genotype and be able to evaluate the MHC information in the signals of conspecifics (Boehm & Zufall 2006, Hettyey et al. 2010). Previously, researchers have shown that condition-dependent signals of quality can be used by both sexes to assess potential mates (Penn & Potts 1998a & 1998b, Charpentier et al. 2010, Johansson & Jones 2007). Although evidence of correlation with MHC genotype has derived primarily from visual signals, such as antler size (Ditchkoff et al. 2001) or bright coloration (Setchell et al. 2009), chemical signals could prove more reliable for advertising MHC genotype (Blaustein 1981, Penn 2002, Mays & Hill 2004, Ziegler et al. 2005): Notably, because degraded MHC molecules are shed from the cell surface and found in body fluids (e.g., serum, saliva, sweat, urine, and glandular secretions), they may function directly as odorants (Singh et al. 1987, Milinski et al. 2005, Boehm & Zufall 2006). MHC molecules may also bind relevant volatile compounds (Leinders-Zufall et al. 2004, Willse et al. 2005, Aksenov et al. 2012, but see Kwak et al. 2010). Lastly, the MHC may influence the composition of the host’s microbiota (Lanyon et al. 2007, Zomer et al. 2009, Archie & Theis 2011), including those dwelling within scent glands that contribute to volatile chemical production (Gorman et al. 1974, Theis et al. 2013, Leclaire et al. 2017a & 2017b). Among taxa that display MHC-associated mate choice, researchers have implicated the operation of an olfactory mechanism in fish (Reusch et al. 2001, Aeschlimann et al. 2003, Milinski et al. 2005), reptiles (Olsson et al. 2004), birds (Ekblom et al. 2004, Leclaire et al. 2014), and mammals (Yamazaki et al. 1976, Radwan et al. 2008), including humans (reviewed in Havlicek & Roberts 2009, Winternitz et al. 2016).

In terms of a model species for an odor-based test of MHC signaling (e.g., Knapp et al. 2006) and discrimination of mate quality, ring-tailed lemurs are ideal because of their elaborate system of olfactory communication (Jolly 1966). Animals of both sexes engage in genital scent marking, depositing chemically complex secretions that share ∼170 volatile compounds (Scordato et al. 2007). The diversity and relative abundance of these chemicals contain information about the signaler’s sex, breeding condition, individual identity, and genome-wide microsatellite diversity (or neutral heterozygosity), as well as its relatedness to other individuals (Scordato et al. 2007, Charpentier et al. 2008, Boulet et al. 2009 & 2010, Crawford et al. 2011). Moreover, this chemically encoded information is salient and distinguishable to conspecifics (Scordato & Drea 2007, Charpentier et al. 2010, Crawford et al. 2011). Thus, lemur genital odors honestly advertise at least one measure of genetic quality and relatedness in both sexes. To test if ring-tailed lemurs also advertise their MHC quality and dissimilarity via olfactory signals, we genotyped animals at the most diverse MHC gene, DRB, analyzed the volatile chemical composition of their genital secretions, and used behavioral testing to determine if conspecifics can use genital scent to discriminate between MHC genotypes, according to either the good genes or good fit models.

## METHODS

### Subjects

Our subjects (*N* = 62) derived from three captive populations of ring-tailed lemurs, located at the Duke Lemur Center (DLC, *N* = 24 females and 24 males) in Durham, NC, the Indianapolis Zoo (*N* = 8 females and 4 males) in Indianapolis, IN, and the Cincinnati Zoo (*N* = 2 females) in Cincinnati, OH. All of the animals were healthy adults that were reproductively intact (i.e., neither gonadectomized nor hormonally contracepted) at the time of study. They were housed in mixed-sex pairs or groups, with similar living conditions and provisioning routines across all three institutions (for more details, see Scordato et al. 2007). Animal care met with institutional guidelines and was in accordance with regulations of the United States Department of Agriculture. The research protocols were approved by the Institutional Animal Care and Use Committee of Duke University (protocol numbers A245-03-07 & A143-12-05) and by the research directors of each zoo.

As described below, although all individuals were genotyped for MHC-DRB diversity, not all individuals participated as secretion donors for chemical analyses or bioassay presentation, nor did all individuals participate as bioassay recipients. To achieve appropriate sample sizes while working with an endangered species often requires years of sample collection and observation, which presents various logistical challenges (for more details, see Drea et al. 2013). Specific to our work here, secretion sampling is restricted because of several factors, including but not limited to facility practices which limit captures in order to avoid undue stress for the animals, and unavailability of an animal due to hormonal contraception (Crawford et al. 2011), pregnancy (Crawford & Drea 2015), mortality, or transfer between facilities. Participation as a bioassay recipient may also be precluded because of participation in previous studies leading to a lack of any ‘unknown’ donor odors to present. For this study, however, we have overcome these challenges wherever possible through long-term study or the addition of animals from other facilities for use as participants.

### MHC genotyping

Using DNA extracted from whole blood or tissue, we genotyped the subjects at the MHC-DRB loci using parallel tagged next-generation sequencing (Grogan et al. 2016). Briefly, we amplified a 171-bp fragment, excluding primers, of the 270-bp second exon of the MHC-DRB gene. This fragment is the most frequently genotyped MHC loci in non-model primate species, especially in lemur species for which genomic data to design primers are scarce (e.g., Schwensow et al. 2007, Huchard et al. 2012, Sommer et al. 2013, Pechouskova et al. 2015, Kaesler et al. 2017). Because this fragment excludes several variable amino acids within the MHC-DRB gene, we are likely underestimating the amount of MHC-DRB variability present. Nonetheless, because the genotyped fragment represents the most variable part of exon 2, we can use this 171-bp fragment as a proxy of diversity across the 6 exons of MHC-DRB. We sequenced pooled amplicons using parallel tagged sequencings on two platforms: Ion Torrent PGM^®^ 314v2 chips (Life Technologies, Grand Island, NY, USA) and 454 Titanium^®^ 1/8th lanes (Roche, Nutley, NJ, USA). True MHC-DRB alleles were distinguished from artefacts using a published workflow (Grogan et al. 2016). Each ring-tailed lemur possessed a mean ± S.D. of 2.22 ± 0.92 MHC-DRB alleles (range = 1-4; see Supplemental Table 1, adapted from Grogan et al. 2017). Next, because of the degeneracy of the genetic code and the similarity in the physiochemical properties of some amino acids, we organized these MHC-DRB alleles (*N* = 20) into MHC-DRB ‘supertypes’ (*N* = 13; Grogan et al. 2017). Supertypes are groups of MHC alleles that have similar antigen binding properties despite having different nucleotide sequences (Doytchinova & Flower 2005, Schwensow et al. 2007), and thus are likely to bind the same subset of pathogen peptides. Supertypes are determined first by aligning the MHC-DRB sequences with the human HLA-DRB sequence (Brown et al. 1993) to identify antigen binding sites. Then, amino acid sites under positive selection are identified using the CODEML analysis in PAML (Version 4.7; Yang 2007). For amino acids identified as under putative positive selection, we imported their physiochemical properties, including hydrophobicity, steric bulk, polarity, and electronic effects (Sandberg et al. 1998), into a matrix in Genesis 1.7.6 (Sturn et al. 2002). Finally, using hierarchical clustering via Euclidean distance methods, we identified supertypes based on antigen binding similarity. The range of the number of alleles collapsed into each supertype grouping is 1-8 alleles, with a mean ± S.D. of 2.01 ± 1.54 alleles.

**Table 1.** Explanatory variables used in the analysis of recipient responses to conspecific odorants and the model under which they were tested.

| Variable | Model | Definition |
| --- | --- | --- |
| MHC<sub>supertype diff</sub> | Good Fit | The number of MHC-DRB supertypes that are different between a recipient-donor dyad |
| MHC<sub>allele diff</sub> | Good Fit | The number of MHC-DRB alleles that are different between a recipient-donor dyad |
| MHC<sub>min</sub> | Good Fit | The minimum number of amino acid differences between a recipient-donor dyad's most similar MHC-DRB allele |
| MHC<sub>mean</sub> | Good Fit | The mean number of pairwise amino acid differences between a recipient-donor dyad |
| MHC<sub>aa diff</sub> | Good Fit | The total number of amino acid differences between all possible pairs of alleles between a recipient-donor dyad |
| MHC<sub>donor</sub> | Good Genes | The number of MHC-DRB alleles present in the donor |

### Genital secretion sample collection

We collected secretion samples from the genital glands of ‘donor’ animals from two captive ‘populations’: the DLC over 10 years (2003-2013, *N* = 24 females, 24 males) during the breeding and nonbreeding seasons, and the Indianapolis Zoo (2011, *N* = 8 females, 1 male) during the breeding season. For ring-tailed lemurs in the Northern Hemisphere, we considered samples collected from November to March to be ‘breeding season’ samples and those collected from May to August to be ‘nonbreeding season’ samples (Drea 2007, Scordato et al. 2007). At the DLC, trained handlers carefully caught and gently restrained the animals, which were awake and habituated to these procedures. At the Indianapolis Zoo, collections occurred during annual physical examinations, performed by Zoo staff members, while the animals were under anesthesia. No secretions were collected from subjects at the Cincinnati Zoo. Following published methods (Scordato et al. 2007), we used pre-cleaned cotton swabs and forceps to collect the secretions. We placed the scented swabs in pre-cleaned chromatography vials and stored the vials at −80 °C until their use in chemical analysis or behavioral experiments. We have previously shown that individual-specific scent signatures are stable across both years and storage time (Scordato et al. 2007, Crawford et al. 2011, Drea et al. 2013). These odors were used for either chemical analyses or bioassay presentation, based upon the season of collection, the number of odors available per individual, and the number of possible recipients to which the odor could be presented. In order to maximize the possible bioassay presentations, we prioritized achieving an appropriate sample size for chemical analyses to detect statistical differences rather than analyzing the chemistry of every individual.

### Gas chromatography-mass spectrometry

All chemical analyses were performed on secretions collected from subjects at the DLC. We used previously published methods to quantify the volatile chemical composition of a subset of these genital secretions (collected from *N* = 20 females, 23 males). These data include previously published data from some individuals (*N* = 17 females, 19 males; Scordato et al. 2007, Charpentier et al. 2008, Boulet et al. 2010), as well as unpublished data from additional individuals (N = 3 females, 5 males). Briefly, we extracted the volatile components of the secretions into 1.5 ml of methyl-*tert*-butyl ether, concentrated the extraction, and analyzed the components on a Shimadzu GCMS-QP2010 instrument (Shimadzu Scientific Instruments) equipped with a Shimadzu AOC-20 series autosampler. The compounds were detected using the automatic peak detector (SOLUTION WORKSTATION software, Shimadzu Scientific Instruments) and the peaks individually verified via consultation with the National Institute of Standards and Technology library (for further details, see Drea et al. 2013).

For analyses of the chemical data, we discarded compounds with inconsistent retention times or that did not comprise at least 0.05% of the overall area of the GC-MS chromatogram. The remaining compounds (*N* = 338 compounds from female genital secretions and 203 compounds from male genital secretions) consisted of fatty acids, fatty acid esters, cholesterol derivatives, alkanes, and other unidentified compounds (Scordato et al. 2007, Charpentier et al. 2008, Boulet et al. 2010). To represent the overall chemical composition of lemur genital secretions (Charpentier et al. 2008), we used three measures of diversity: richness, the Shannon index, and the Simpson index (Legendre & Legendre 1998, McCune et al. 2002). For our purposes, richness is the absolute number of compounds present per chromatogram regardless of relative abundance or rarity. By contrast, the Shannon and Simpson diversity indices reflect the relative abundances in different ways: The Shannon index is primarily influenced by common compounds of intermediate abundance, whereas the Simpson index gives more weight to compounds of the greatest relative abundance (McCune et al. 2002, Charpentier et al. 2008). We calculated these diversity indices for each individual’s overall chemical profile, as well for specific subsets of compounds that have been implicated in signaling fertility: fatty acids (*N* = 33/338 and 25/203 in female and male genital secretions, respectively) and fatty acid esters (*N* = 112/338 and 87/203; Boulet et al. 2010, Crawford et al. 2011).

### Behavioral bioassays

To test if ring-tailed lemur ‘recipients’ can use the secretions of ‘donors’ to discriminate between the MHC genotypes of opposite-sex conspecifics, we conducted 300 behavioral trials or ‘bioassays,’ (Scordato et al. 2007, Charpentier et al. 2008, Greene et al. 2016). To maximize our use of ‘unknown’ donors per recipient (i.e., the recipient had never resided in a group with the donor nor had smelled secretions from the donor during a previous bioassay experiment), we used recipients (*N* = 27) from the multiple institutions, including at the DLC (*N* = 5 females and 14 males), Cincinnati Zoo (*N* = 2 females), and Indianapolis Zoo (*N* = 2 females and 4 males), and secretion samples from ‘unknown’ donors of the opposite sex at the DLC (*N* = 31 females and 18 males). Bioassays were conducted from November-February in 2011 and 2012 following previously established protocols (see Scordato et al. 2007, Charpentier et al. 2008, Greene et al. 2016). First, we temporarily isolated a recipient from its group members, then secured three fresh wooden dowels in a row to the fence of the animal’s enclosure (separated by 20 cm) at a 45° angle to the ground. The center dowel served as an unscented control, whereas a ∼2 cm area of the outer dowels (at lemur nose level) were rubbed for ∼10-15 seconds with the scented swabs from different donors, to simulate a naturally placed scent mark. Each recipient underwent 1-3 trials per day over 4-6 days, with each trial lasting 10 minutes, ultimately participating in 8-12 trials in total.

We presented the secretions to each recipient in randomized order. We also maximiated the number of donor dyads whose secretions could be presented across recipients, while minimizing the number of times we presented a given donor’s secretions to any recipient (average ± S.D. exposures = 1.85 ± 1.05, range = 0-6). Upon completion of the day’s trials, the recipient was reunited with its group.

The bioassays were videotaped and the videos were scored by three observers, using an established ethogram. Prior to scoring experimental trials, we calculated inter-observer reliability from five ‘practice’ trials. Differences in the labeling of an event or in the chronology or timing (>1 s) were considered disagreements (Martin & Bateson 2007, Scordato et al. 2007) and scoring of videos did not commence until inter-observer reliability scores exceeded 90%. The behavior recorded included the recipient’s proximity to each dowel and its sniffing, licking, biting, genital marking and, for males only, shoulder rubbing, wrist marking, and tail rubbing (Supplemental Table 2, adapted from Scordato et al. 2007). We also recorded where investigatory or scent-marking behavior occurred relative to each scent ‘mark’ (i.e., whether the behavior was directed at the mark itself, adjacent to the mark, but on the dowel, or within 15 cm of the dowel). Specifically, we recorded the location of countermarks because their placement could have particular significance: Overmarking or placing one’s mark directly on top of the original mark might mask the original mark, whereas adjacent-marking or placing one’s mark near the original mark leaves the original mark intact (Drea 2014).

**Table 2.** Relationships between measures of chemical diversity and of MHC diversity in male ring-tailed lemurs across seasons, with significant relationships indicated in bold.

| Measure of chemical diversity | Best-fit explanatory variable | Z value | P value | Effect |
| --- | --- | --- | --- | --- |
| Richness-overall | Season | 2.37 | **0.018** | Chemical richness decreases from nonbreeding season to breeding season, but increases with increasing MHC diversity. |
|  | MHC<sub>supertype</sub> | 2.31 | **0.021** |  |
|  | Season * MHC<sub>supertype</sub> | -0.71 | 0.475 |  |
| Shannon Index-overall | Season | 2.43 | **0.015** | Shannon diversity decreases from nonbreeding season to breeding season, but increases with increasing MHC diversity. |
|  | MHC<sub>supertype</sub> | 2.52 | **0.012** |  |
|  | Season * MHC<sub>supertype</sub> | -1.45 | 0.146 |  |
| Simpson Index-overall | Season | 1.64 | 0.10 | Simpson diversity does not change from breeding to nonbreeding season; however, Simpson diversity increases with increasing MHC diversity. |
|  | MHC<sub>supertype</sub> | 2.17 | **0.03** |  |
|  | Season * MHC<sub>supertype</sub> | -0.89 | 0.37 |  |

### Statistical Analyses

#### General analytical procedures

To examine the relationships between MHC-DRB genotype and olfactory ornamentation, as well as the ability of ring-tailed lemurs to discriminate MHC genotype via genital secretion, we analyzed the data in a series of generalized linear mixed models (GLMMs), using the package ‘glmmADMB’ (Version 0.7.7) in RStudio (Version 3.0.2; RStudio 2015). MHC diversity can be measured in several ways (e.g. the number of alleles in an individual’s genotype, the number of supertypes, or the number of amino acid differences between all alleles in an individual’s genotype, described below), as can MHC similarity between individuals. We assessed several measures of MHC diversity for an individual, as well as several measures of similarity between dyads. Because we expected all measures of MHC genetic diversity to be correlated, we evaluated each of the explanatory genetic variables independently, and used Akaike information criteria (AIC) values to determine the best model (Zuur et al. 2009). We considered the model with the lowest AIC value to be the best fit for the data and report only those models. To examine if MHC similarity were reflected in chemical similarity, we used partial Mantel tests to compare the number of un-shared or unique MHC-DRB supertypes to the relative Euclidean distance matrices between male-male (MM), female-female (FF), and male-female (MF) dyads.

#### Analyses of MHC-DRB diversity and chemical complexity in individual males and females

To examine the relationship between MHC-DRB diversity and chemical complexity, we analyzed the sexes separately and each chemical diversity index was evaluated in a separate series of GLMMs using either a Gaussian or gamma distribution (see Supplemental Table 3). We included donor identity as a random variable. Explanatory variables included season (i.e., breeding and nonbreeding) and one of four different genetic variables. Our four measures of MHC-DRB diversity were 1) the absolute number of MHC-DRB alleles (MHC_allele_), 2) the number of MHC-DRB supertypes (MHC_supertype_), 3) the average nucleotide base-pair differences between all alleles within an individual donor (MHC_seqmea__n_), and 4) the mean number of amino acid differences between the alleles possessed by an individual donor (MHC_aamean_; Supplemental Table 3). Because of the skew in frequency of specific MHC-DRB supertypes, i.e., seven supertypes were found in fewer than five individuals whereas one supertype was found in more than 85% of individuals, we were unable to examine if the possession of a specific supertype can be signaled via the chemical complexity of genital secretions. For females, we also analyzed the diversity of two subsets of chemicals, fatty acids (FAs) that are known to signal fertility in female primates (Michael et al. 1974, Doty et al. 1975, Matsumoto-Oda et al. 2003), and fatty acid esters (FAEs), which are synthesized from FAs (Cheng & Russell 2004, Hargrove et al. 2004). We have shown that both chemical subsets are correlated with microsatellite diversity of female ring-tailed lemurs during the breeding season (Boulet et al. 2010). We were unable to control for neutral heterozygosity estimated via microsatellites (see Charpentier et al. 2008, Boulet et al. 2010 for microsatellite methods), because microsatellite data were unavailable for > 20% of our subjects. Nevertheless, we did assess the correlation between microsatellite heterozygosity and MHC diversity, i.e., the number of MHC-DRB alleles and MHC-DRB supertypes. Using the subset of subjects for which both genetic measures of diversity were available (*N* = 36), we found no correlation between microsatellite heterozygosity and the number of MHC-DRB alleles within an individual (correlation coefficient = 1.056, F = 1.51, *P* = 0.227), and no correlation between microsatellite heterozygosity and the number of MHC-DRB supertypes (correlation coefficient = −0.8812, F = 1.461, *P* = 0.235; however, see Grogan et al. 2017). As we had previously found no correlation between individual chemical diversity and adult age, month of collection within season, or DLC housing condition (Charpentier et al. 2008), we did not include these co-variables in our analyses. Lastly, to ensure that the few individuals with the most diverse MHC-DRB genotype were not driving any association between MHC-DRB diversity and chemical diversity, we re-ran the final GLMMs after removing the most diverse individuals from both the male (*N* = 1) and female analyses (*N* = 1).

**Table 3.** Relationships between measures of chemical diversity and MHC diversity in female ring-tailed lemurs across seasons, with significant relationships indicated in bold.

| Measure of chemical diversity | Best-fit explanatory variable | Z value | P value | Effect |
| --- | --- | --- | --- | --- |
| Richness-overall | Season | -0.53 | *0.600* | No relationship between female richness and season or MHC diversity. |
|  | MHC<sub>supertype</sub> | -1.02 | *0.310* |  |
|  | Season * MHC<sub>supertype</sub> | -0.92 | *0.360* |  |
| Shannon Index-overall | Season | 0.77 | *0.442* | Overall Shannon index possibly negatively correlated with MHC diversity, but only in the nonbreeding season. |
|  | MHC<sub>supertype</sub> | -0.34 | *0.733* |  |
|  | Season * MHC<sub>supertype</sub> | -1.69 | *0.091* |  |
| Simpson Index-overall | Season | 0.86 | *0.390* | No relationship between female Simpson index and season or MHC diversity. |
|  | MHC<sub>supertype</sub> | 0.24 | *0.810* |  |
|  | Season * MHC<sub>supertype</sub> | -0.98 | *0.330* |  |
| Shannon Index - FAs | Season | 1.62 | *0.104* | FA Shannon index negatively correlated with MHC diversity, but only in the nonbreeding season. |
|  | MHC<sub>supertype</sub> | -0.20 | *0.838* |  |
|  | Season * MHC<sub>supertype</sub> | -3.09 | **0.002** |  |
| Shannon Index - FAEs | Season | 1.43 | *0.152* | FAE Shannon index negatively correlated with MHC diversity, but only in the nonbreeding season. |
|  | MHC<sub>supertype</sub> | -0.42 | *0.678* |  |
|  | Season * MHC<sub>supertype</sub> | -2.63 | **0.009** |  |

#### Analysis of MHC relatedness and chemical similarity between all possible dyads

To investigate if chemical similarity between dyads reflects similarity in their MHC genotypes, we calculated matrices of genetic distances using the number of different MHC supertypes between each dyad, or the number of different putative antigen binding pocket configurations that are present in one individual but not the other. We then estimated chemical distances between pairs of individuals by analyzing all chemicals present in secretion profiles of males (*N* = 203 compounds; Charpentier et al. 2008), females (*N* = 338; Boulet et al. 2010), or in both sexes (*N* = 170; Boulet et al 2009). We calculated relative Euclidean distance matrices for same-sex (MM or FF) and mixed-sex (MF) dyads, respectively, using PC-ORD (version 7.0, McCune & Mefford 2016), and following published protocols (Charpentier et al. 2008, Boulet et al. 2009). We calculated matrices separately as follows: breeding season (22 males, *N* = 231 MM dyads; 17 females, *N* = 136 FF dyads; 39 males and females, *N* = 374 MF dyads); nonbreeding season (20 males, *N* = 190 MM dyads; 18 females, *N* = 153 FF dyads; 38 males and females, *N* = 360 MF dyads). Because MM and FF matrices were square, we assessed linear relationships between chemical and MHC distances using partial Mantel tests in FSTAT (version 2.9.3.2, with 10,000 randomizations; Goudet 2001). As in previous studies (Charpentier et al. 2008, Boulet et al. 2010), we controlled for potentially confounding covariates, including the subject’s age, social housing condition, and the month of secretion sample collection. For the MF comparisons, we first generated full matrices using all possible MM, FF, and MF pairs (breeding season *N* = 704 dyads, nonbreeding season *N* = 741 dyads). We then extracted chemical, genetic, and covariate information for MF dyads only. Unlike MM and FF matrices, the MF matrix was not square. Therefore, we assessed relationships with 10,000 Spearman’s correlation permutation tests using the JMUOUTLIER package in R (Version 1.3; Garren 2016), as in the study by Slade et al. (2016). Lastly, we assessed the correlation between MHC similarity within dyads (i.e., the number of unique or unshared MHC-DRB alleles and supertypes between two individuals) with dyad relatedness, as measured by the Queller and Goodnight index (IDQG calculated by Boulet et al. 2010). We found that although dyad relatedness was significantly correlated with MHC dissimilarity for both number of MHC-DRB alleles (*N* = 629 dyads, slope = −0.71, T-value = −4.21, P = 0.000029) and number of MHC-DRB supertypes (*N* = 629 dyads, slope = −0.67, T-value = −4.23, *P* = 0.000027), the negative relationships explained less than 3% of the variance in either correlation (R^2^ = 0.026 and R^2^ = 0.026, respectively).

#### Behavioral analyses of mixed-sex, recipient-donor combinations

We explored the relationship between the recipients’ behavioral responses to donor secretions and measures of absolute and relative MHC-DRB diversity between the mixed-sex, recipient-donor dyads (Table 1). To test for patterns that would be consistent with disassortative mating under the good fit model, we used several measures of dissimilarity and sequence divergence between each recipient-donor dyad (Schwensow et al. 2008, Setchell et al. 2010, Huchard et al. 2013). We also used the donor’s number of MHC alleles (MHC_donor_) to examine if, under the good genes model, the secretion of potential ‘mates’ with the greatest MHC diversity were distinguished, regardless of their dissimilarity. Lastly, we also examined non-linear relationships between MHC diversity and dissimilarity between dyads by including the quadratic forms of all genetic explanatory variables in our GLMMs. Quadratic terms were retained only if the AIC value was better than the GLMM that included only linear terms.

In our analyses, we excluded all recipient behavior that occurred in < 5% of trials and any behavior that did not show a significant differential response via Wilcoxon signed-rank tests in favor of the test dowels over the blank, control dowel (Supplemental Table 4; Charpentier et al. 2010). Ultimately, for male recipients, we analyzed proximity to the pole, sniffing and licking the mark and surrounding areas, shoulder rubs, and wrist marking the area adjacent to the mark. For females, we analyzed sniffing the mark and the adjacent area, as well as licking the mark. In each GLMM, we also included the non-genetic explanatory variables of trial number on a given day (i.e., 1-3), the number of times that a recipient had been presented with the secretion of a given donor over the course of the study (i.e., 1-6), as well as the secretion donor nested under secretion recipient, as a random term. For each behavioral response, we used either a Poisson or negative binomial distribution depending on the dispersion of the data (Supplemental Table 5) and applied a zero-inflation correction factor if appropriate. Initially, each GLMM was constructed using all explanatory terms, including both a linear and quadratic term for the genetic variable to account for potential nonlinear relationships. The GLMM with the lowest AIC value was chosen for further exploration, for which we sequentially dropped explanatory terms, then compared these models using AIC values. We report only the best fit models for each response variable below.

**Table 4.** Partial Mantel tests for same-sex (MM and FF) dyads, showing seasonal relationships between genital odor distance (relative Euclidean) and MHC-based genetic distance in ring-tailed lemurs. Odor distance is based on 203 and 338 compounds for MM and FF dyads, respectively. Tests include three socio-demographic and environmental variables as covariates. Significant (*P* ≤ 0.05) sums of squares (SS), partial Mantel correlation coefficient (*r*), and *P* values are shown in bold type, and trending (*P* ≤ 0.10) values are shown in italics.

| Dyad type | Variable | Number of unique MHC supertypes |  |  |  |  |  |
| --- | --- | --- | --- | --- | --- | --- | --- |
|  |  | Breeding season |  |  | Nonbreeding season |  |  |
| | | SS | $r$ | $P$ | SS | $r$ | $P$ |
| MM dyads | MHC | **2.309** | **0.408** | **<0.001** | 0.042 | -0.079 | 0.276 |
|  | Age | 0.203 | 0.121 | 0.068 | **0.139** | **0.145** | **0.005** |
|  | Housing | 0.157 | 0.106 | 0.101 | 0.006 | -0.029 | 0.692 |
|  | Month of collection | 0.006 | -0.020 | 0.763 | *0.124* | *0.137* | *0.062* |
| FF dyads | MHC | **0.391** | **0.313** | **<0.001** | 0.003 | 0.027 | 0.738 |
|  | Age | 0.001 | 0.014 | 0.872 | 0.008 | -0.047 | 0.560 |
|  | Housing | 0.029 | 0.085 | 0.323 | 0.004 | 0.034 | 0.679 |
|  | Month of collection | *0.093* | *0.153* | *0.076* | **1.461** | **0.623** | **<0.001** |

**Table 5.** Relationship between the MHC-DRB genotype of female odorant donors and male recipient behavior, directed toward the female’s scent mark, with significant relationships indicated in bold. Explanatory variables with superscripts indicate the quadratic variable, while those without superscripts are linear.

| Behavior | Best-fit explanatory variable | Z value | P value | Effect |
| --- | --- | --- | --- | --- |
| Sniff mark | MHC<sub>supertype diff</sub><sup>2</sup> | -2.10 | **0.036** | Shorter duration with increasing supertype differences between dyad |
| Sniff pole | MHC<sub>supertype diff</sub><sup>2</sup> | -2.54 | **0.011** | Shorter duration with increasing supertype differences between dyad |
| Lick mark | MHC<sub>donor</sub> | -4.45 | **<0.001** | Shorter duration with increasing female donor MHC diversity |
|  | MHC<sub>donor</sub><sup>2</sup> | 4.14 | **<0.001** |  |
| Shoulder rub | MHC<sub>supertype diff</sub><sup>2</sup> | -2.09 | **0.037** | Fewer rubs with increasing supertype differences |

## RESULTS

### Signaling of individual quality via odor-gene covariance

We found that both sexes of ring-tailed lemurs signaled individual MHC quality via genital secretion chemistry, although the correlates of signaling differed between males and females: Male MHC diversity was positively correlated with chemical diversity, regardless of season (Figure 1; Table 2). Removal of the most diverse male from the analyses still resulted in significant relationships between MHC-DRB diversity and Shannon diversity (*P* = 0.011), between MHC-DRB and season (*P* = 0.007), and a trend toward significance for the interaction between MHC-DRB diversity, Shannon diversity, and season (*P* = 0.068). By contrast, female MHC diversity was unrelated to overall chemical diversity in either season (Figure 2; Table 3); nevertheless, it was negatively correlated to the diversity of two important subsets of chemicals, FAs and FAEs, during the nonbreeding season. Similarly, removal of the most diverse female from the analysis of overall Shannon diversity and MHC-DRB diversity results in no significant relationships. After removal of the same female from the analyses of the chemical subsets of fatty acids and fatty acid esters, the interaction between season and MHC-DRB diversity in relation to Shannon diversity of fatty acids is no longer statistically significant although still trending in the same direction (*P* = 0.10). The relationship between fatty acid ester Shannon index in females and season retains a trending relationship (*P* = 0.053) and MHC-DRB diversity (*P* = 0.067), although their interaction is not significant (*P* = 0.161).

**Figure 1.**
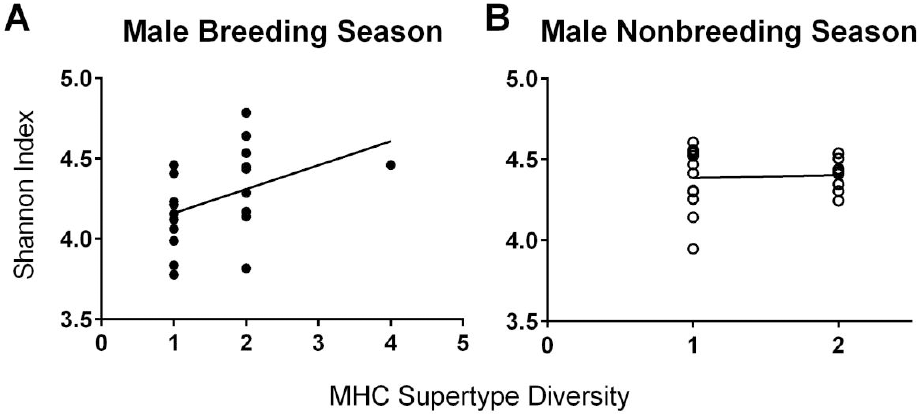
Seasonal relationships between chemical diversity in genital secretions and MHC supertype diversity in male ring-tailed lemurs. Overall chemical diversity increased with increasing MHC supertype diversity across the breeding season (A), shown in black circles, and the nonbreeding season (B), shown in white circles.

**Figure 2.**
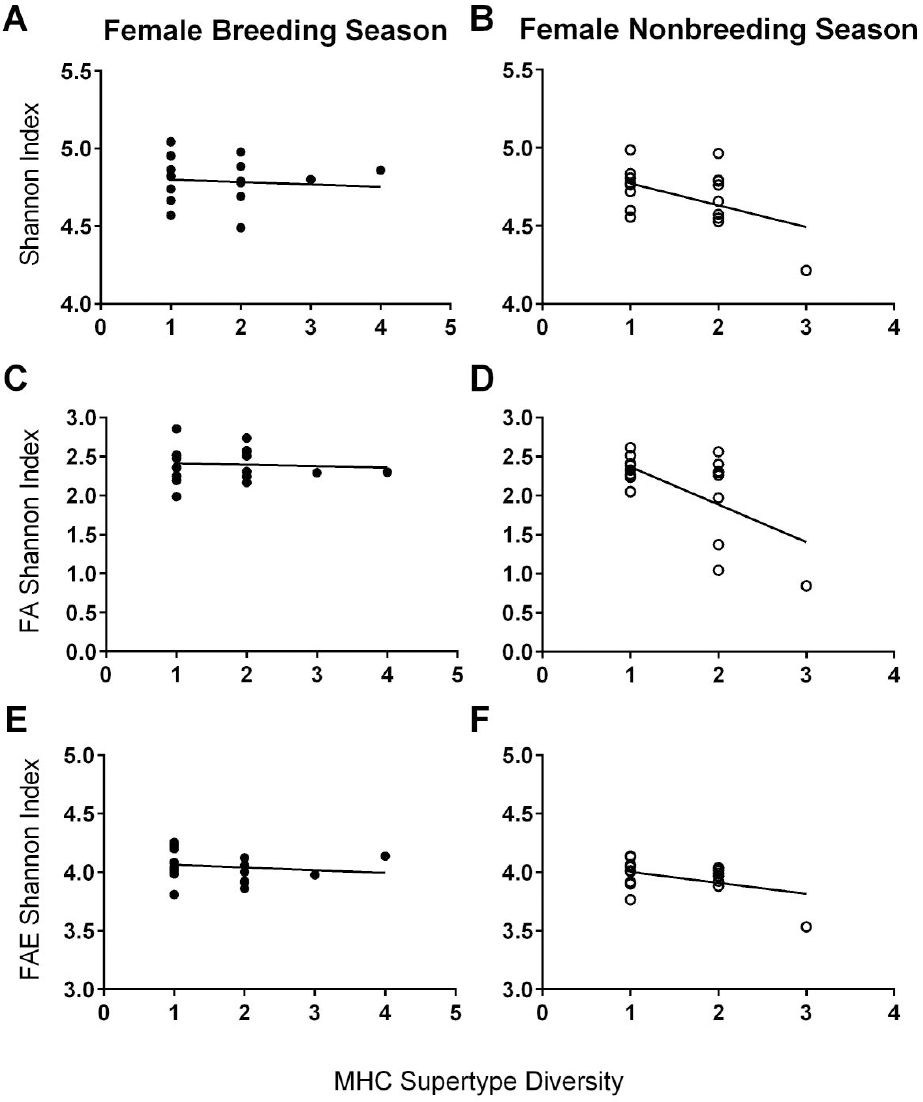
Seasonal relationships between various measures of chemical diversity in genital secretions and MHC supertype diversity in female ring-tailed lemurs. Overall chemical diversity was not correlated with MHC supertype diversity in either the breeding season (A), shown in black circles, or nonbreeding season (B), shown in white circles. Diversity indices for fatty acids (FAs) and fatty acid esters (FAEs) were not correlated with MHC supertype diversity in the breeding season (C and E, respectively), but were negatively correlated in the nonbreeding season (D and F, respectively).

### Signaling of relatedness via dyadic, odor-gene covariance

In all same-sex lemur dyads, genital olfactory cues encoded information about MHC distance, but in a season-dependent fashion (Figure 3; Table 4). After controlling for covariates, chemical distances between MM dyads correlated with unique MHC supertypes during the breeding season (*P* < 0.001, Figure 3A-B), but not during the nonbreeding season (*P* = 0.270, Figure 3C-D). Similarly, for FF dyads, we observed significant, season-specific correlations between the number of unique MHC supertypes and chemical distances (breeding season *P* < 0.001, Figures 3E-F; nonbreeding season *P* = 0.729, Figure 3G-H).

**Figure 3.**
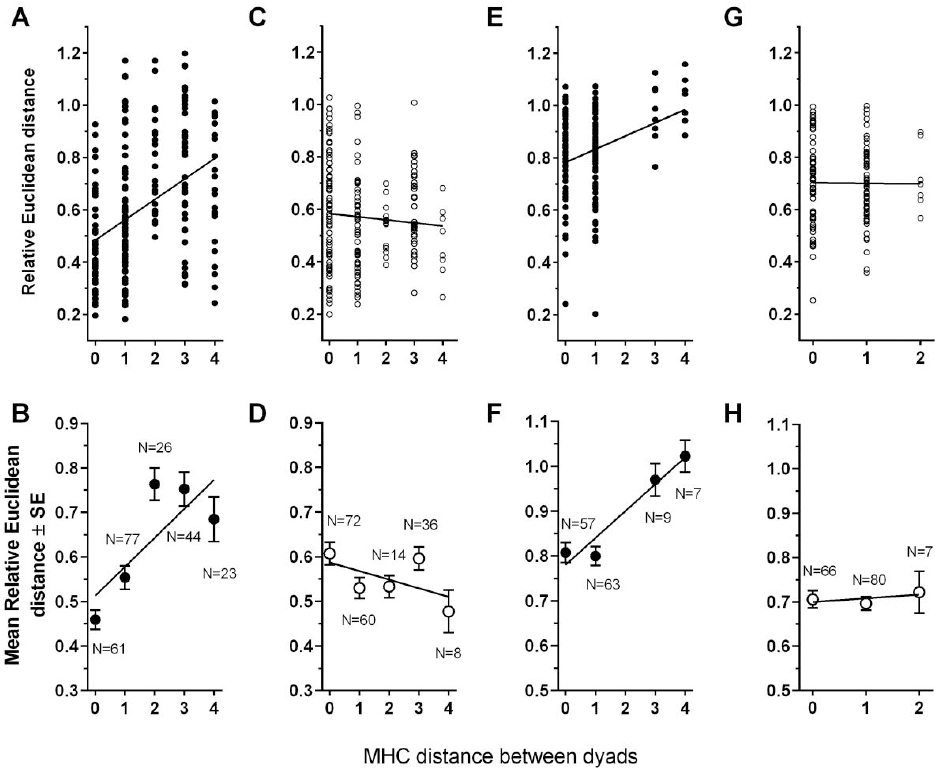
Relationships between genetic distance (number of unique MHC supertypes, i.e., MHC_supertype diff_) and chemical distance (relative Euclidean) between male-male (A-D) and female-female (E-H) ring-tailed lemur dyads during the breeding (closed circles; A-B and E-F) and nonbreeding seasons (open circles; C-D and G-H). The numbers of dyads are provided in the bottom panel.

Although we could not find any relationship between chemical distance and MHC distance between MF dyads in the breeding season (*r* = 0.0014, *P* = 0.8280; Figure 4A), but we did find a trending negative relationship between mixed-sex dyads during the nonbreeding season (*r* = − 0.0099, *P* = 0.0647; Figure 4B).

**Figure 4.**
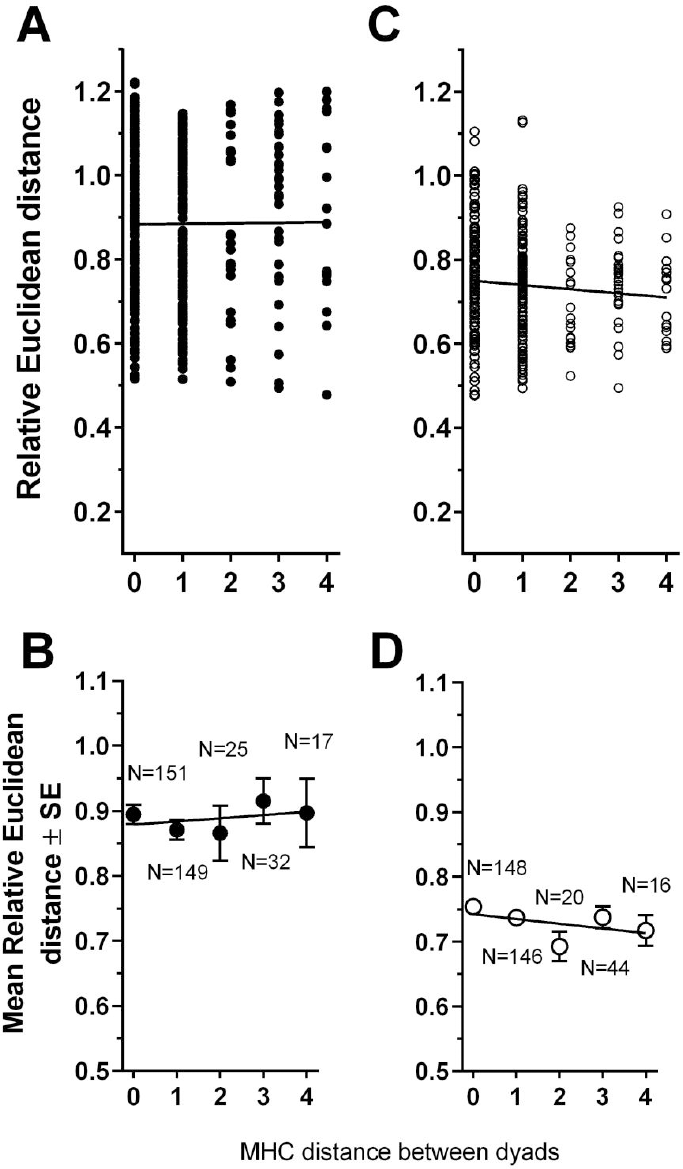
Relationships between the number of unique MHC supertypes (i.e., MHC_supertype diff_) and chemical distance (relative Euclidean) between male-female, ring-tailed lemur dyads during the breeding season (A-B), shown in black circles, and nonbreeding season (C-D), shown in white circles. The numbers of dyads are provided in the bottom panel.

### Olfactory discrimination of MHC genotype between mixed-sex conspecifics

Although in the breeding season, we could only detect the chemical signaling of MHC diversity in males, both male (Table 5; Figure 5) and female (Figure 6) recipients showed significant behavioral discrimination between the genital secretions of opposite-sex, conspecific donors based on their possession of different MHC-DRB genotypes. The pattern of response to conspecific secretions, however, differed between the sexes.

**Figure 5.**
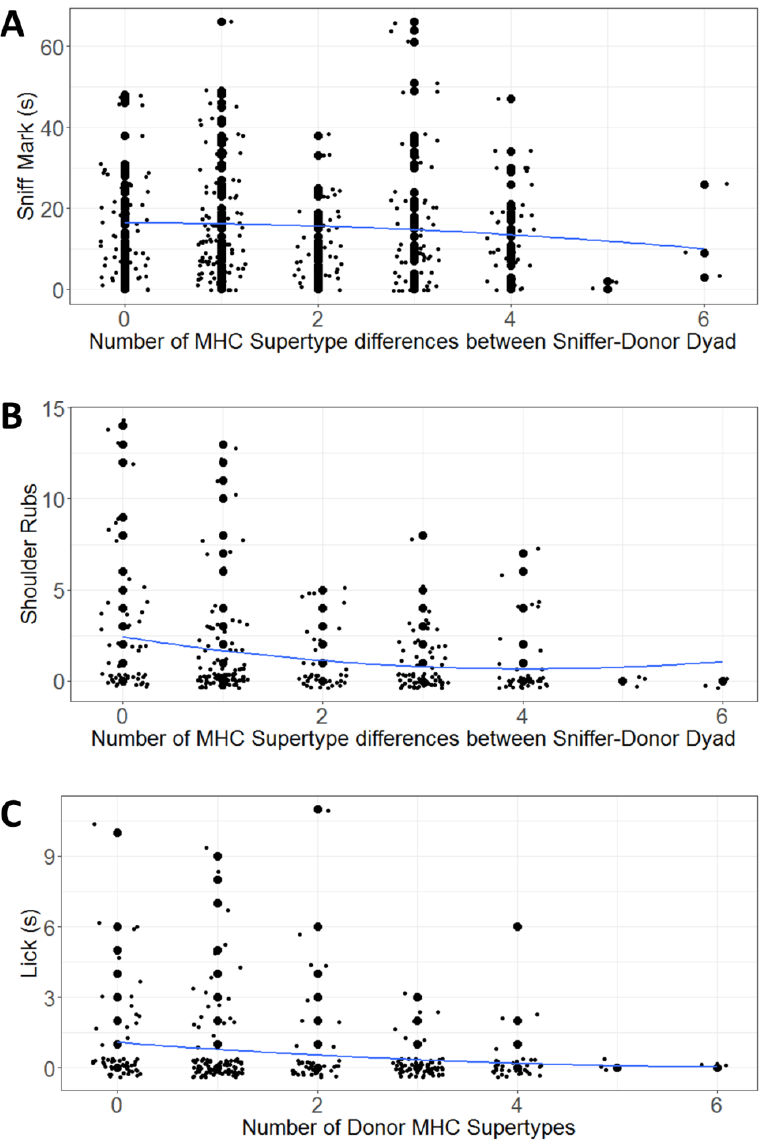
Response of male ring-tailed lemurs to the odorants of female conspecifics: As the number of MHC-DRB supertypes between the male recipient and the female donor increases, the duration of male sniffing (A) and the frequency of shoulder rubbing (B) decreases significantly. As the diversity of the MHC genotype of the female donor increases, the duration of licking (C) decreases. Points are jittered to avoid overlap of data.

**Figure 6.**
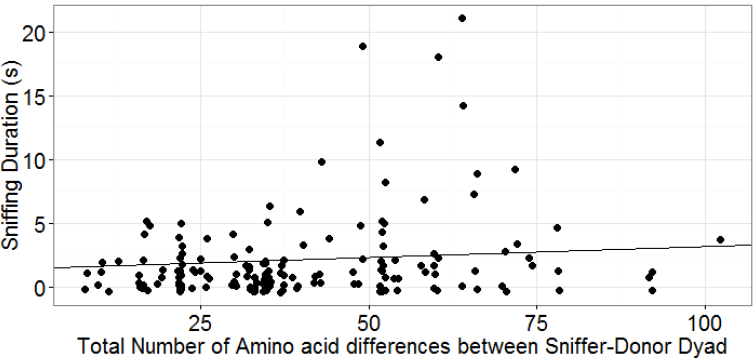
Response of female ring-tailed lemurs to the odorants of male conspecifics: As the number of amino acid differences between female recipient and male odorant donor increases, the duration of time the female sniffs the male’s odor increases.

Male recipients did not differ in the time they spent in proximity to either of the scented dowels, nor did they differ in the number of wrist marks they directed to the area adjacent the mark; however, they investigated female secretions more if the donors were more similar at the MHC-DRB to themselves than if the donors were more dissimilar at the MHC-DRB. Specifically, compared to their responses to the secretions of MHC-DRB dissimilar females, male recipients spent more time sniffing the secretions of MHC-DRB similar females, more time sniffing the area adjacent to the mark, and responded with more shoulder rubs (Table 5; Figure 5).

Additionally, as the absolute MHC-DRB diversity of female donors increased, male recipients also licked the mark more frequently (Table 5). Because the frequency of occurrence and duration of behavior were correlated for licking, shoulder rubbing, and wrist marking, we report on only one measure (either frequency or duration) per behavior (Table 5; Figure 5).

Female recipient responses to the mark itself did not differ according to the MHC-DRB diversity of male donors; nevertheless, female recipients did direct differential responses towards the areas adjacent to the male’s mark. Specifically, as the MHC-DRB amino acid dissimilarity of the male donor increased, female recipients spent more time sniffing areas adjacent to the mark (z value = 3.11, *P* = 0.002; Figure 6).

## DISCUSSION

Owing to its role in survival and reproductive success, immunogenetic diversity is an important predictor of mate quality and preference and may be signaled via visual or chemical means. Our study provides support for the socially salient, chemical signaling of genetic quality in a strepsirrhine primate, as would be necessary according to the good genes and good fit paradigm for choosing a mate or social partner. Despite sex differences in the chemical ‘indicators’ of quality and their seasonal emergence, ring-tailed lemurs of both sexes signaled their MHC diversity and dissimilarity to conspecifics via the volatile component of their genital secretions. Moreover, both sexes were able to use these and potentially other olfactory cues to discriminate relevant information about the MHC genotypes of opposite-sex conspecifics. These results confirm the functional significance of our previous work showing detectable relationships between chemical diversity and microsatellite diversity in both sexes (Charpentier et al. 2008, Boulet et al. 2010, Charpentier et al. 2010). Our results also provide a foundation from which to explore if ring-tailed lemurs actually choose mates according to the good genes and/or good fit models using reproductive success data from wild populations.

More specifically, males appeared to signal both MHC-DRB diversity and microsatellite diversity, or their good genes, in a similar fashion: Both measures of genetic diversity were positively correlated with overall chemical diversity of genital secretions, although the relationship with microsatellite diversity only emerged in the breeding season, whereas the MHC odor-gene covariance emerged regardless of season (albeit more strongly in the breeding season). Females, however, signaled their good genes, i.e., genetic diversity, via specific chemicals, e.g., fatty acids and fatty acid esters. Previously, we had shown that females signaled increased microsatellite diversity via a negative relationship with the diversity of fatty acids during the breeding season. Here, we show that females signal MHC diversity via a negative relationship with FA and FAE diversity, but in the nonbreeding season instead.

Potentially, females may be signaling different information depending on the season. During the breeding season, signaling genome-wide microsatellite diversity and relatedness may be critical to avoid inbreeding (Boulet et al. 2009, 2010). In contrast, signaling MHC-specific diversity and health during the nonbreeding season might convey health and competitive ability during periods of intense female-female competition (e.g., López-Idiáquez et al. 2016). The months of pregnancy and lactation are energetically expensive for female ring-tailed lemurs (O’Mara & Hickey 2014). Additionally, intragroup female competition for access to food increases (Sauther 1993, Gould et al. 2003). During these social disputes, the killing of vulnerable infants is a significant risk, and is committed by both sexes (Hood 1994, Jolly et al. 2000, Ichino 2005, Charpentier & Drea 2013, Kittler & Dietzel 2016). Signaling one’s health and vitality may reduce the likelihood of aggressive encounters that could lead to infanticide by competing females (reviewed in Stockley & Campbell 2013).

Our results contrast the lack of odor-gene covariance found in mandrills (*Mandrillus sphinx*), the only other primate in which a relationship between secretion chemistry and MHC diversity has been investigated. In both male and female mandrills, MHC diversity was unrelated with the chemical diversity of secretions obtained from the surface of the sternal gland (Setchell et al. 2011). MHC information, however, may be signaled through aspects of their olfactory signature that were not analyzed by the authors. For instance, just as female ring-tailed lemurs signal MHC and microsatellite diversity through a subset of chemicals (i.e., fatty acids and fatty acid esters; Boulet et al. 2010), so too might MHC-DRB information be contained in the composition and relative abundance of a subset of the compounds present in certain secretions. Alternatively, information may be encoded in the non-volatile portion of the secretion (Hurst et al. 2001, Beynon et al. 2002, Brennan & Zufall 2006, Kwatra & Drea 2007) or be signaled through the composition of the microbiota present in the gland (Boehm & Zufall 2006, Zomer et al. 2009, Archie & Theis 2011, Pearce et al. 2017) and the odorants they produce (Gorman et al. 1974, Theis et al. 2013). Further exploration of individual compounds, specific subsets of chemicals, or the non-volatile fraction of the mandrill secretion might yield a signaling pattern that conveys information about MHC genotype. Such evidence would support findings that male mandrills appear to use the MHC genotype of a potential mate for mate guarding decisions (Setchell et al. 2016) and that MHC diversity is correlated with male reproductive success (Setchell et al. 2010).

The chemical composition of lemur genital secretions also signals MHC dissimilarity, or fit, between male-male, female-female, and male-female dyads, echoing previous results demonstrating the same pattern for microsatellite diversity (Charpentier et al. 2008, Boulet et al. 2009 & 2010). Signaling relatedness to any potential social ‘partner’ could be relevant throughout the year, to avoid related competitors or beneficially direct nepotism (Charpentier et al. 2008, 2010). Signaling relatedness or compatibility to opposite sex conspecifics would be particularly important in the breeding season to avoid inbreeding and maximize offspring diversity (Brown 1999, Tregenza & Wedell 2000, Neff & Pitcher 2005). Evidence now exists that odorants signal MHC dissimilarity within same-sex and opposite-sex dyads in two taxa formerly thought to be primarily visually oriented, namely birds (black-legged kittiwake: Leclaire et al. 2014; song sparrows: Slade et al. 2016) and anthropoid primates (mandrills: Setchell et al. 2011), suggesting greater relevance of olfactory cues than previously suspected.

Regarding behavior, our male recipients responded most to the scent of females that had the most similar MHC-DRB genotypes to their own. Such a pattern may reflect ‘preferences,’ indicating that males may be avoiding outbreeding depression (Sommer 2005b), seemingly contrary to the good fit prediction that individuals should choose mates to maximize their dissimilarity. Nevertheless, an increase in response time may not necessarily indicate an increase in interest. Instead, increased male investigation could simply reflect that more processing time was required to decipher the female’s potential as a mate. Previously, in a study of microsatellite diversity, we had shown that male ring-tailed lemurs spent more time sniffing the secretions of less-related females (Charpentier et al. 2010). Regardless of the potential contradiction, both sets of findings indicate that male ring-tailed lemurs are minimally able to discriminate conspecifics according to both overall genetic relatedness and MHC diversity/dissimilarity. Lastly, our finding that female ring-tailed lemurs spent the most time sniffing the vicinity of secretions from MHC-DRB dissimilar males complements previous work showing that females of other species show greater response to the scents of more MHC dissimilar males than of more MHC-diverse males.

We have confirmed an honest olfactory mechanism of ornamentation and potential mate choice, namely via genital odor-MHC gene covariance and discrimination, in both sexes of ring-tailed lemurs. Likewise, information about immunogenetic quality and similarity may also influence general social behavior, specifically for prioritizing agonistic or nepotistic interactions. Female lemurs are expected to be choosy under the traditional paradigm of sexual selection (Trivers 1972); however, mate choice may be equally important for male ring-tailed lemurs (Parga 2006). In this species, females socially dominate their male conspecifics (Jolly 1966), are strictly seasonal and generally fertile only 1-3 times per year for a period of less than 24 hours (Evans & Goy 1968, Van Horn & Resko 1977), and often cycle somewhat synchronously with other females in the social group (Pereira 1991). Thus, both sexes should be choosy about the competitive effort directed towards their potential partners. Our data extend the potential for olfactory-based MHC discrimination across the primate order and add to a growing body of literature suggesting that mate choice may depend on both MHC dissimilarity and diversity (Kamiya et al. 2014).

## Acknowledgements

This work was supported by the National Science Foundation (BCS #0409367, IOS #0719003) to CMD, the Duke University Center for Science Education to KEG, and Duke University to CMD and KEG. We would like to thank the staff and animal technicians of the Duke Lemur Center, the Indianapolis Zoo, and the Cincinnati Zoo for their assistance during sample collection and bioassay trials. We are especially grateful to Erin Ehmke, David Brewer, and Britt Keith at the Duke Lemur Center, to Lynn Villers, Robert Shumaker, and Holly Blaylock at the Indianapolis Zoo and to Terri Roth and Ronald Evans at the Cincinnati Zoo. We could not have accomplished this study without them. We would also like to thank Jillian Wisse for her help with conducting the bioassays, as well as Laura Damiani and Bernice Kwan for their help in scoring the videos. This manuscript is DLC publication #XXXX.

## Author Contributions

CMD and KEG conceived of the idea for this study with help from MB & RLH. KEG produced the MHC-DRB genotypes and conducted the behavioral bioassays, and MB & RLH produced the secretion chemistry data. KEG, MB, & RLH performed the analyses and KEG wrote the original draft of the manuscript. CMD critically revised the manuscript with assistance from KEG, RLH, and MB. All authors have approved the final manuscript for publication.

## Data Accessibility

CSV files of MHC, chemical, and behavioral data from bioassays, will be deposited into Dryad. Analyses reported in this article can be reproduced using these data.

## Conflict of Interest

The authors declare no conflict of interest.

